# Partnership between epigenetic reader BRD4 and transcription factor CEBPD

**DOI:** 10.1101/2020.03.27.012674

**Authors:** Qingwei Wang, Mengxue Zhang, Go Urabe, Bowen Wang, Hatice Gulcin Ozer, Yitao Huang, K. Craig Kent, Lian-Wang Guo

## Abstract

Vascular smooth muscle cell (SMC) state/phenotype transitions underlie neointimal hyperplasia (IH) predisposing to cardiovascular diseases. Bromodomain protein BRD4 is a histone acetylation reader and enhancer mark that co-activates transcription elongation. CCAAT enhancer binding protein delta (CEBPD) is a transcription factor typically studied in adipogenesis and immune cell differentiation. Here we investigated the association between BRD4 and CEBPD in SMC state transition.

Chromatin immunoprecipitation sequencing (ChIPseq) showed enrichment of BRD4 and histone acetylation (H3K27ac) at *Cebpd* and enhancer in rat carotid arteries undergoing IH. In vitro, BRD4 silencing with siRNA reduced SMC expression of CEBPD. Bromodomain-1 but not bromodoamin-2 accounted for this BRD4 function. Endogenous BRD4 co-IP’ed with CEBPD; *Cebpd* promoter and enhancer DNA fragments co-IP’ed with CEBPD or endogenous BRD4 (ChIP-qPCR). These co-IPs were abolished by the BRD4 bromodomain blocker JQ1. TNFα upregulated both BRD4 and CEBPD. Silencing CEBPD averted TNFα-induced inflammatory SMC state transition (heightened IL-1β, IL6, and MCP-1 mRNA levels), so did JQ1. CEBPD overexpression increased PDGFRα preferentially over PDGFRβ; so did TNFα, and JQ1 abolished TNFα’s effect.

Our data reveal a BRD4/CEBPD partnership that promotes CEBPD’s own transcription and inflammatory SMC state transition, thus shedding new light on epigenetic reader and transcription factor cooperative actions in SMC pathobiology.

## Introduction

Smooth muscle cells (SMCs) as a major component and signaling hub maintain vascular wall homeostasis. However, upon extra- and intra-cellular perturbations, they undergo various state transitions, losing innate identity and function while acquiring new phenotypes. This SMC plasticity is now known as enabled by epigenetics^1^ – regulations critical in biology particularly in development and disease. Depending on micro-environmental cues, SMCs may transition to inflammatory and/or migro-proliferative or other states, contributing to a range of vascular disorders including neointimal hyperplasia (IH) that obstructs circulation^2^. Therefore, in the pursuit of new interventional paradigms, it is important to interpret the IH pathogenic mechanisms from an epigenetic perspective. However, epigenetic determinants of SMC state transitions only begin to be unveiled^2,3^.

Our previous reports suggested BRD4 as a determinant of SMC state transitions in vitro and IH in vivo^4, 5^, motivating further research to decipher its molecular underpinning. BRD4 belongs to the family of BETs (Bromo/ExtraTerminal domain-containing proteins), including BRD2, BRD3, BRD4, and BRDT (testis-restricted). Rapid progress, particularly in cancer research, reveals that histone acetylation reader BRD4 is closely associated with cell state/identity changes and hence critical to an array of pathogenic processes^6^. As a result, BRD4 is being intensively targeted in human trials, for cancers and beyond. BRD4 is found to potently prompt RNA polymerase II pause release thereby co-activating transcription elongation in response to stimulation^6^. This action involves multiple factors such as the elongation factor complex and central machinery of transcription. It remains poorly understood how BRD4, a seeming global regulator, assumes its functional specificity in different cell types and micro-environments^2^. Transcription factors (TFs) and enhancers are thought to confer BRD4 some gene loci specificity in the genomic landscape^6, 7^. However, little is known as to how BRD4 and specific TFs and enhancers govern different SMC state transitions^8^.

The CCAAT enhancer binding proteins (CEBPs) comprise a family of TFs highly versatile in their pathophysiological functions. They are most notably involved in adipogenesis and immune cell differentiation among other activities. CEBPA and CEBPB are best studied, whereas CEBPD is much less understood. Evidence collectively implicates CEBPD as a central player in responses to inflammatory stimuli^9^, as mostly reported in immune cells with a paucity of data from SMCs. CEBPD’s function is highly contextual. Dichotomous or even opposing effects mediated by CEBPD have been reported, e.g. in cell proliferation and in macrophage differentiation^9^. Obviously, CEBPD’s functional regulations reported in other cell types or signaling contexts cannot be simply extrapolated to SMCs. Interestingly, while CEBPD expression is generally low in normal conditions, it rapidly increases in response to environmental perturbations, such as arterial injury that induces IH^10, 11^. This raises an important question as to what epigenetic factors govern CEBPD’s level and functional impact on SMC state transition^12^.

In the current study, as we sought to identify BRD4’s downstream functional mediators and CEBPD’s upstream epigenetic determinants in SMCs, the two searches converged. Through chromatin immunoprecipitation sequencing (ChIPseq), we observed that angioplasty injury induced enrichment of BRD4 and H3K27ac at the CEBPD gene. Indeed, BRD4 dominated CEBPD expression in SMCs. Moreover, CEBPD co-immunoprecipitated with BRD4, with its own promoter, and with BRD4/H3K27ac-associated enhancer. The cooperativity between BRD4 and CEBPD manifested in heightened SMC expression of pro-inflammatory cytokines. On the whole, this study explored a previously unknown area, where we found a BRD4/CEBPD partnership that promoted the inflammatory SMC state transition.

## Results

### Angioplasty injury induces BRD4 and H3K27ac enrichment at the CEBPD gene in rat carotid arteries

We previously found that blocking BETs’ bromodomains with JQ1 effectively curbed neointima progression in an authentic model of angioplasty-induced IH in rat carotid arteries^4, 13^. While BRD4 drastically increased in the angioplasty-injured artery wall, in vitro and in vivo data suggested that BRD4 is a determinant of IH^4, 5, 13^. In an effort to further identify the mediators of BRD4’s pro-IH function, we performed high-throughput ChIPseq with injured (and uninjured) carotid arteries using the same angioplasty model of IH. Angioplasty abruptly alters the SMC micro-environment by endothelium denudation and mechanical damage of the artery wall. This exposes SMCs to a myriad of blood-borne stimulants typical of cytokines such as TNFα and growth factors including PDGF, which potently trigger various SMC state transitions that perpetuate IH. Arteries were collected for ChIPseq at post-angioplasty day 7 as this is the peak time of pro-IH molecular and cellular events^14, 15^. Epigenomic marks involved in transcriptional activation, including BRD4, H3K27ac, and H3K4me1^16^, were chosen for ChIP experiments, and the main body of data is available in our other report^17^. BRD4 and H3K27ac are associated with active enhancers, and H3K4me1 can be found with active, inactive, or poised enhancers.

We noticed that the ChIPseq intensity peaks for BRD4 and H3K27ac enriched at the gene of CEBPD, a TF poorly studied for SMC state transitions and IH. Indicating genomic co-localization of BRD4, H3K27ac, and H3K4me1, their ChIPseq peaks aligned in the same genomic region (Figure 1A). Moreover, Venn diagram (Figure 1B) shows highly overlapping genes annotated based on the genomic DNA fragments that co-IP’ed with BRD4 and H3K27ac. By contrast, ChIPseq signal for H3K27me3 (a gene repression mark) was negligible. Together, these data suggest that BRD4 may regulate the expression of transcription factor CEBPD in balloon-injured rat carotid arteries.

**Figure 1.**
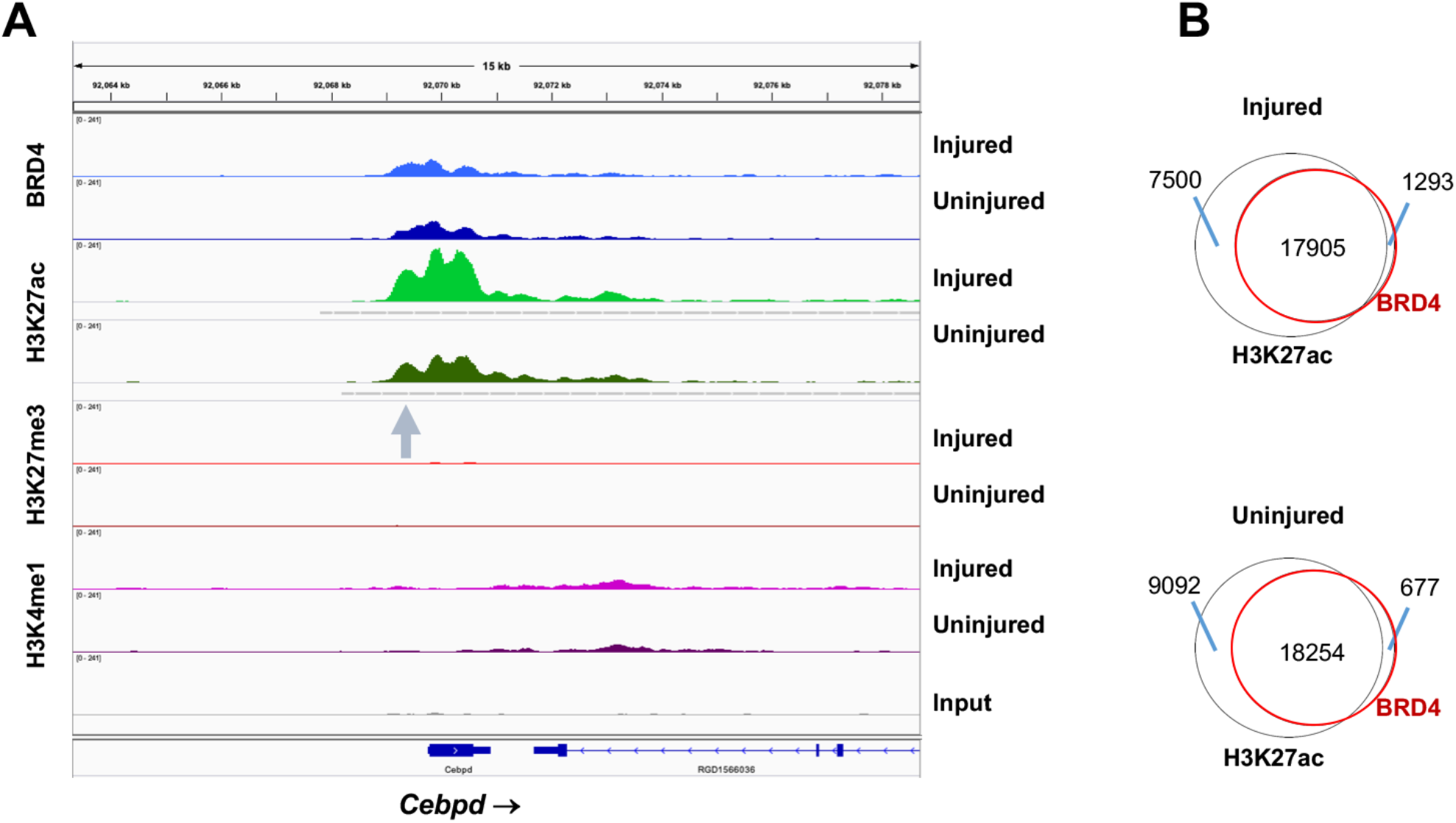
Injury-induced enrichment of BRD4 and H3K27ac at Cebpd in rat carotid arteries. Balloon-injured rat left common carotid arteries and contralateral arteries (uninjured control) were collected at day 7 post angioplasty and snap frozen until use for ChIPseq. **A.** ChIPseq binding density at *Cebod.* Shown are ChIPseq peak profiles for BRD4 and histone marks. Data were obtained from injured and uninjured arteries. Non-specific input indicates low background noise. **B.** Venn diagrams showing overlap of ChIPseq peaks for H3K27ac and BRD4.

### BRD4 but not BRD2 silencing diminishes CEBPD expression, so does BRD4 dominant-negative bromodomain-1

To distinguish whether BRD4 is the only BET family member that determines CEBPD expression, genetic silencing was achieved in MOVAS (an established cell line of SMCs) using siRNAs specific to each of the 3 BETs. As shown in Figure 2A, while each siRNA was effective in silencing its respective target BET, BRD4 silencing, but not BRD2 or BRD3 silencing, led to pronounced reduction of CEBPD mRNA. This result was confirmed by Western blots, where only BRD4 silencing reduced CEBPD protein (Figure 2, B and C). BRD4 interacts with specific chromatin loci by reading/binding histone acetyl-lysine bookmarks through its two tandem bromodomains (BD1 and BD2), which are therefore crucial for BRD4’s epigenetic functions. We thus dissected their role in regulating CEBPD expression. Interestingly, expressing a dominant-negative BD1 domain (i.e. a competitor to endogenous BD1) substantially reduced CEBPD protein, whereas a dominant-negative BD2 domain that proved functional in our other report^5^ did not have an effect herein (Figure 3, A-C). Consistently, treating SMCs with Olinone, an inhibitor selective to BD1 in BETs^19^, diminished CEBPD protein levels, whereas RVX208 (selective to BD2 in BETs)^20^ did not significantly alter CEBPD expression (Figure 3, D-F). Therefore, our data distinguished that BRD4 but not BRD2 or BRD3 was the determinant BET that controlled CEBPD expression. Furthermore, BD1 rather than BD2 was responsible for this BRD4 function; this result was unexpected given that in previous reports BD2 *vs* BD1 was shown to play a major role in non-SMC settings^5, 21^.

**Figure 2.**
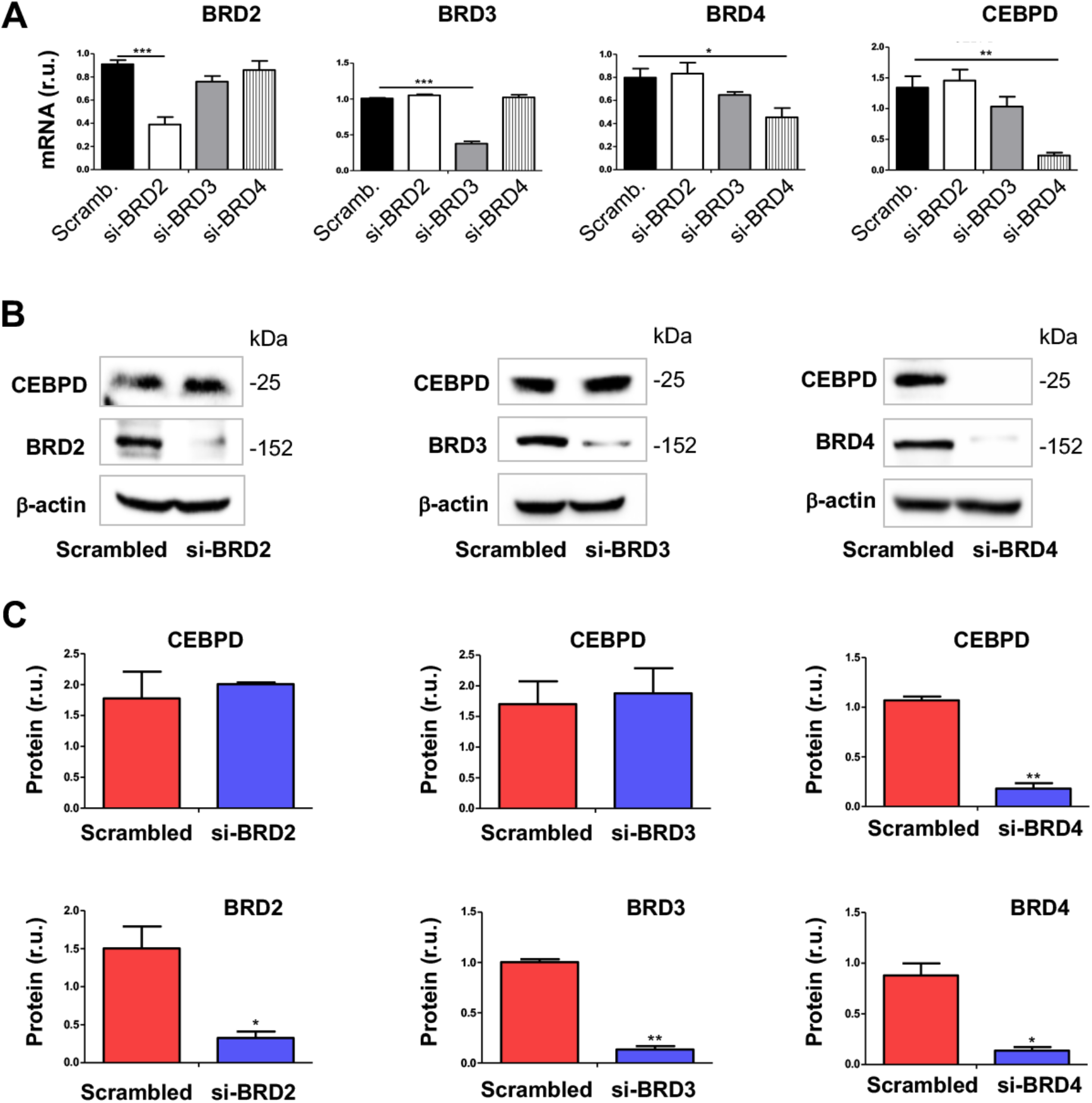
Silencing BRD4 reduces CEBPD mRNA and protein in SMCs. MOVAS cells were transfected with scrambled or specific siRNA for 24h, cultured with fresh medium (no RNAi Max) for another 24-48h, and then harvested for qRT-PCR (A) or Western blot (B and C) analysis, respectively. Quantification: Readings from triplicate qRT-PCR reactions were normalized (to GAPDH) and averaged. The average values from 3 independent repeat experiments were then averaged again to calculate mean ± SEM (n =3). Densitometry of Western blots from independent repeat experiments was normalized (to β-actin, similar band intensity among blots) and then averaged to calculate mean ± SEM (n =3). Statistics: One-way analysis of variance (ANOVA) followed by Bonferroni post-hoc test; *P<0.05, **P<0.01.

**Figure 3.**
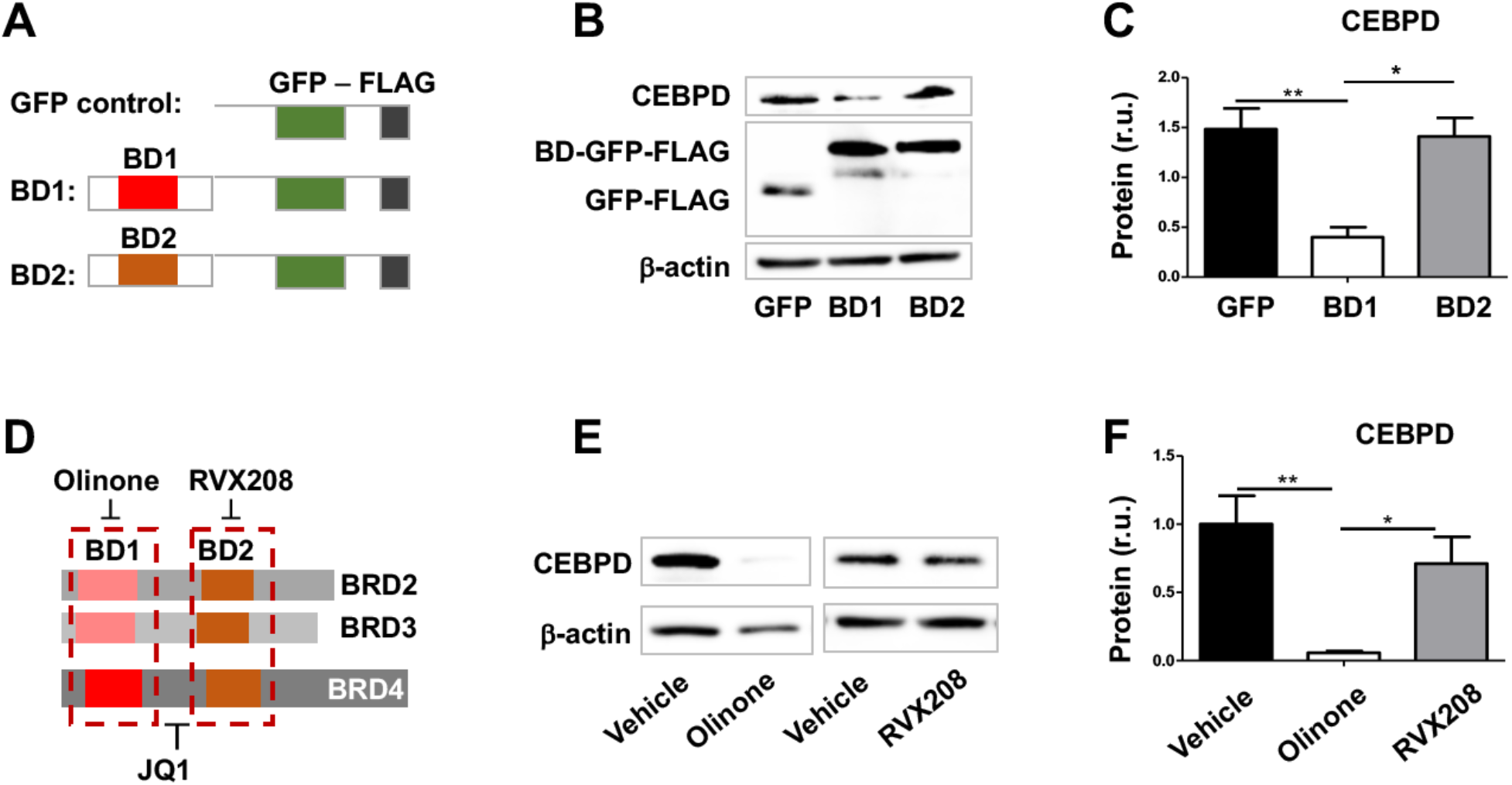
Expression of dominant-negative BD1 of BRD4 inhibits CEBPD expression. **A.** Diagram of the constructs used to exogenously express GFP (control) and a dominant-negative domain competing with the BD1 or BD2 of BRD4. **B** and **C**. Effect of exogenously expressed BD1 or BD2 (of BRD4) on CEBPD expression. **D.** Diagram to show selective binding of olinone and RVX208 to BD1 and BD2, respectively, and binding of JQ1 to both; each of the bromodomain blockers binds all three BETs (BRD2, BRD3, BRD4). **E** and **F**. Effect of bromodomain blockers on CEBPD expression. Data from vehicle blots were pooled into a single value bar in the plot. MOVAS cells were transfected with the dominant-negative BD1 or BD2 plasmid for 24h and cultured in fresh medium (no Lipofectamine) for another 24h before harvest for Western blot analysis. For pharmacological pretreatment, cells were incubated with vehicle (equal amount of DMSO) or a bromodomain blocker for 4h before harvest. Quantification: Densitometry of Western blots from independent repeat experiments was normalized (to β-actin, similar band intensity among blots) and then averaged to calculate mean ± SEM (n =3). Statistics: One-way analysis of variance (ANOVA) followed by Bonferroni post-hoc test; *P<0.05, **P<0.01.

### SMC endogenous BRD4 co-immunoprecipitates with the CEBPD protein

Reports have highlighted the importance of TFs in mediating BRD4’s function of co-activating specific gene expression in cell types other than SMCs. CEBPD is a TF reportedly involved in SMC dysfunction^22, 23^. However, a BRD4/CEBPD cooperativity in SMC pathobiology was yet to be discovered. We therefore performed co-immunoprecipitation (IP) experiments to probe a possible BRD4/CEBPD interaction. As shown in Figure 4, endogenous BRD4 IP’ed with CEBPD specifically (*vs* background control), suggesting their direct interaction, or indirect association but in the same protein complex. Indicative of a role for bromodomains in this interaction, the pan-BETs inhibitor JQ1 which competes with histone acetyl-lysine for binding to BD1 and BD2, blocked the BRD4/CEBPD co-IP. Furthermore, an inhibitor (TSA) of histone deacetylases commonly used to preserve histone acetylation, enhanced the co-IP of BRD4 with CEBPD. These results revealed a previously unidentified BRD4/CEBPD physical association.

**Figure 4.**
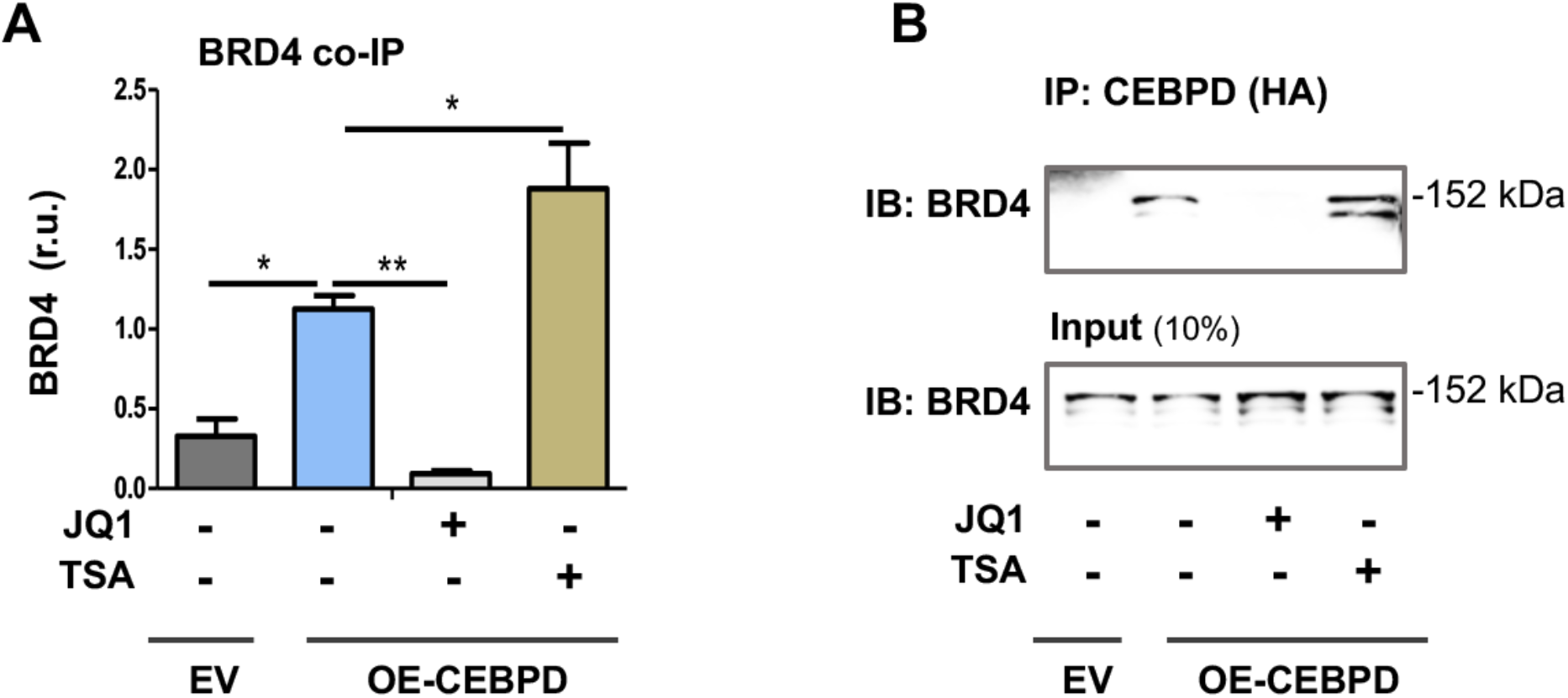
Co-immunoprecipitation of endogenous BRD4 with CEBPD. MOVAS cells stably expressing HA-tagged empty vector (EV) or HA-tagged CEBPD were incubated with vehicle (equal amount of DMSO) or bromodomain blocker JQ1 (1 μM) or HDAC inhibitor TSA (1 μM) for 24h. The cells were then harvested for IP of CEBPD using an antibody against the HA tag. **A.** Quantified data. **B.** Representative Western blots. Quantification: Densitometry of Western blots from independent repeat experiments was normalized (to β-actin, similar band intensity among blots) and then averaged to calculate mean ± SEM (n = 6). Statistics: One-way analysis of variance (ANOVA) followed by Bonferroni post-hoc test; *P<0.05, **P<0.01.

### CEBPD and BRD4 both associate with the Cebpd promoter

To further investigate the observed BRD4/CEBPD protein association in regulating gene expression, we performed ChIP-qPCR experiments. To validate the methodology, we chose the PDGFRα gene promoter for proof-of-principle since PDGFRα is known as a direct target of CEBPD’s TF function in SMCs^22^. As shown in Figure 5A, treatment of SMCs with TNFα stimulated co-IP of PDGFRα promoter DNA with CEBPD. By contrast, the effect on PDGFRβ (previously regarded as non-target^22^) was minor (Figure 5B). CEBPD gain-of-function (overexpression) strongly enhanced PDGFRα promoter DNA pulldown (by ~10-20 fold) either in the presence or absence of TNFα, whereas no effect on PDGFRβ promoter was observed, which could serve as a negative control. Consistent with a BRD4 involvement, pretreating SMCs with JQ1 abrogated PDGFRα promoter pulldown that was enhanced either by TNFα or CEBPD overexpression.

**Figure 5.**
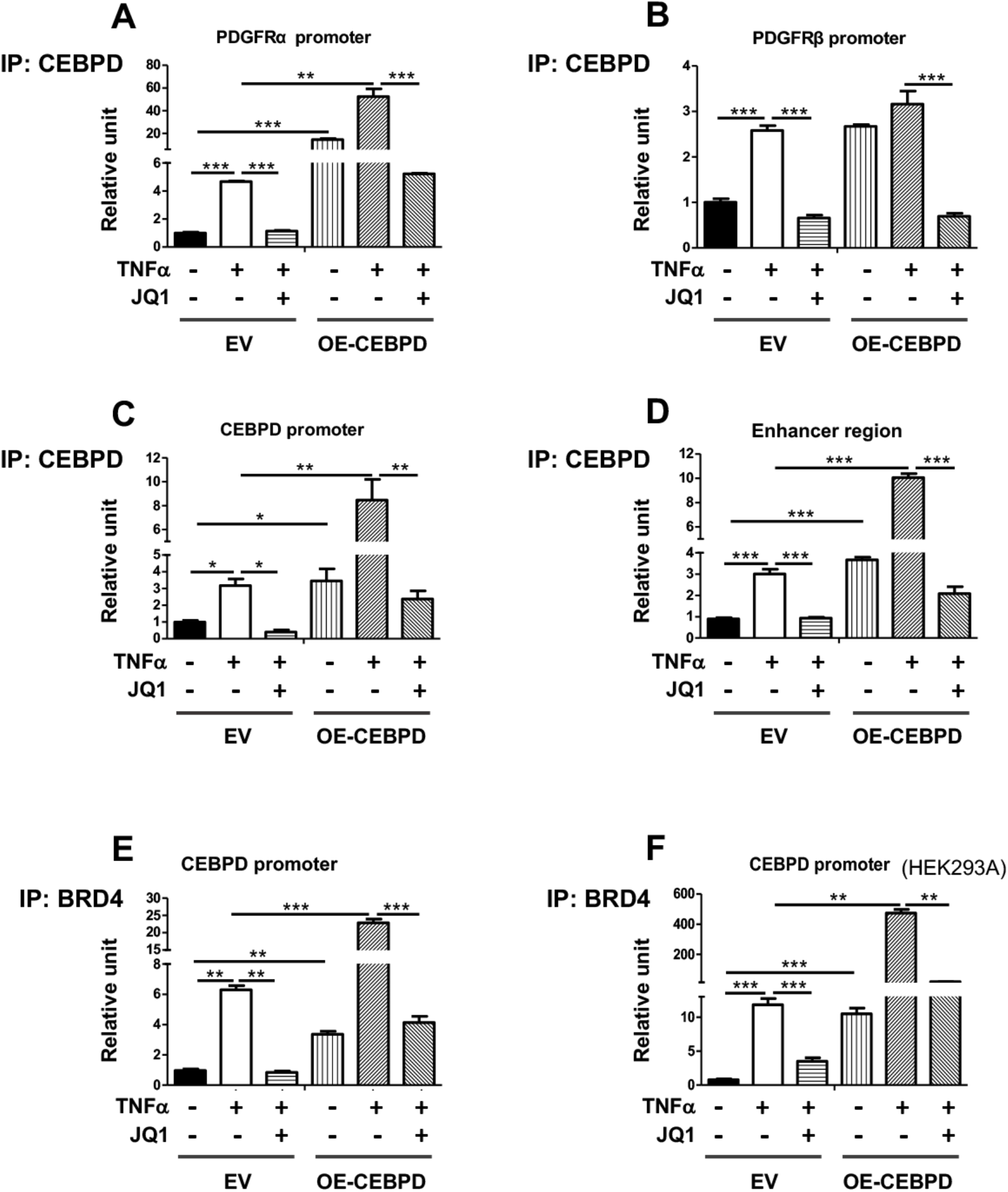
Chromatin immunoprecipitation of Cebpd promoter DNA with CEBPD or endogenous BRD4. **A-D**. ChIP against HA-tagged CEBPD in SMCs. **E-F**. ChIP against endogenous BRD4 in SMCs (E) or HEK293 cells (F). MOVAS cells for stable expression of HA-tagged empty vector (EV) or HA-tagged CEBPD (OE-CEBPD) were starved in basal medium (0.5% FBS) for 24h, pretreated with vehicle or bromodomain blocker JQ1 (1 μM) for 2h, and then treated with TNFα (final 20 ng/ml) for 24h prior to harvest for ChIP-qPCR analysis. Quantification: Readings from triplicate qRT-PCR reactions were normalized (to GAPDH) and averaged. The average values from at least 3 independent repeat experiments were then averaged again to calculate mean ± SEM (n =3-5). Statistics: One-way analysis of variance (ANOVA) followed by Bonferroni post-hoc test; *P<0.05, **P<0.01, ***P<0.001.

More interestingly, software prediction suggested potential CEBPD binding sites on its own promoter. Indeed, ChIP-qPCR indicated that while the co-IP of *Cebpd* promoter DNA with CEBPD increased (by ~2-3 fold) due to TNFα stimulation, CEBPD gain-of-function further markedly enhanced this result (Figure 5C). In either case, JQ1 pretreatment abolished the enhancement of *Cebpd* promoter co-IP with CEBPD, suggesting an essential role of BRD4 in this interaction. BRD4 is an enhancer mark that enriched at *Cebpd,* and inhibiting BRD4 reduced CEBPD expression (Figures 1-3). We therefore inferred that BRD4-dependent enhancer was probably involved in the BRD4/CEBPD complex regulating *Cebpd* transcription. In support of this proposition, we indeed observed that BRD4-dependent enhancer DNA co-IP’ed with CEBPD (Figure 5D), a result highly similar to that of *Cebpd* promoter in all the conditions tested here.

To further confirm an involvement of BRD4 in the transcription-regulating complex, we performed ChIP-qPCR experiments using an antibody to IP endogenous BRD4. We found that while TNFα stimulated the co-IP of *Cebpd* promoter DNA with BRD4 by ~6 fold, CEBPD overexpression further magnified this effect by ~4 fold (Figure 5E). In either case, JQ1 abolished the increase of co-IP. In an effort to gain additional evidence and to expand this interesting finding beyond SMCs, we observed a similar pattern of the endogenous BRD4/*Cebpd* promoter co-IP using the same conditions but HEK293A cells instead of SMCs (Figure 5F).

On the whole, these (Figure 5) and other results (Figures 1-4) provided strong evidence for a multi-component complex involving H3K27ac reader (BRD4), TF (CEBPD), enhancer enriched with BRD4 and H3K27ac, and *Cebpd* promoter DNA, which may govern CEBPD’s own transcription and other target genes.

### CEBPD’s positive role in TNFα-induced inflammatory SMC state transition involves BRD4

Now that the data had suggested a CEBPD (partnering with BRD4) control over its own expression, the next important question was the functional significance of this mechanism in SMC pathobiology. A role for CEBPD in proliferative SMC state transition has been reported^22^. However, IH is a complex process involving multiple SMC state transitions, a prominent one being SMC inflammation which is critically involved in IH-associated conditions such as atherosclerosis. The inflammatory SMC state transition is typically monitored as upregulation of major pro-inflammatory cytokines including IL-1β, IL6, and MCP-1. Both IL-1β and IL6 are pro-IH and atherogenic; so is MCP-1 which is key to recruitment of inflammatory cells (e.g. activated leukocytes) to the vessel wall^24, 25^. As indicated in Figure 6 (A-D), while the CEBPD mRNA in SMCs markedly increased after TNFα stimulation, this treatment also upregulated IL-1β, IL6, and MCP-1 mRNA to a similar extent. Specifically, CEBPD gain (overexpression) and loss (silencing) of function raised and reduced the expression of these cytokines, respectively, in the absence or presence of TNFα. CEBPD’s signaling functions are highly contextual depending on cell type and stimulants^9^. Early evidence showed a pro-inflammatory role for CEBPD in SMCs. This observation, however, was confounded by that activation of peroxisome proliferator-activated receptor (PPAR)-gamma, CEBPD’s downstream target, inhibited CEBPD expression^26^. In this regard, our study contributed an unambiguous result, that is, CEBPD plays a positive role in the specific setting of TNFα-stimulated inflammatory SMC state transition.

**Figure 6.**
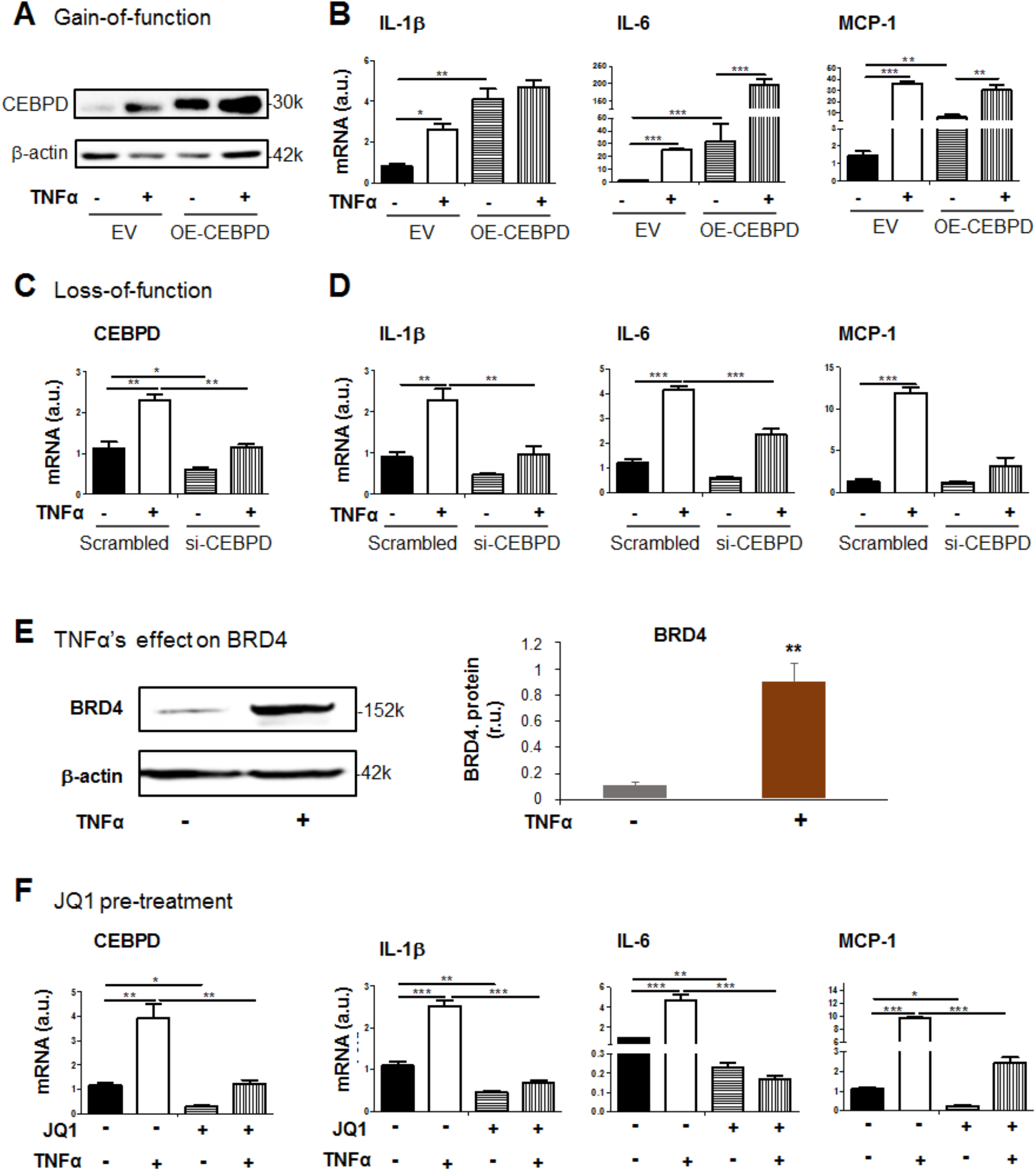
Role of CEBPD in the inflammatory SMC state transition. **A** and **B**. CEBPD gain of function. A shows CEBPD overexpression. **C** and **D**. CEBPD loss of function. C indicates effective CEBPD silencing. **E.** Treatment of SMCs with TNFα upregulates BRD4 protein (Movas cell line). **F.** Effect of pretreatment with bromodomain blocker JQ1. MOVAS cells for stable expression of HA-tagged empty vector (EV) or HA-tagged CEBPD (OE-CEBPD) were starved in basal medium (0.5% FBS) for 24h, pretreated with vehicle or JQ1 (1 μM) for 2h, and then treated with TNFα (final 20 ng/ml) for 24h prior to harvest for qRT-PCR (mRNA) or Western blot (protein) analysis. For CEBPD silencing, MOVAS cells were transfected with siRNA for 24h and cultured in fresh starvation medium (0.5% FBS) for another 24h prior to TNFα treatment. Quantification: Readings from triplicate qRT-PCR reactions were normalized (to GAPDH) and averaged. The average values from at least 3 independent repeat experiments were then averaged again to calculate mean ± SEM (n =3-5). Statistics: One-way analysis of variance (ANOVA) followed by Bonferroni post-hoc test; *P<0.05, **P<0.01, ***P<0.001.

It was interesting to note that TNFα stimulation enhanced the *Cebpd* promoter co-IP with BRD4 (Figure 5E) and consequently elevated CEBPD mRNA (Figure 6C). We therefore determined the effect of TNFα on BRD4 expression, and the data indicated that treating SMCs with TNFα upregulated BRD4 protein (Figure 6E). Since BRD4 rather than BRD2 or BRD3 controlled CEBPD expression via its bromodomain (Figures 2 and 3), we expected that JQ1 which blocks BET bromodomains would reveal BRD4’s function in TNFα-stimulated inflammatory SMC state transition. Indeed, pretreatment with JQ1 abrogated TNFα-stimulated upsurge of the SMC expression of all three cytokines (IL-1β, IL6, MCP-1) (Figure 6F). At this point, the observed BRD4/CEBPD physical association (Figures 4 and 5) appeared to manifest as co-function in regulating SMC inflammation, which was not previously addressed.

### CEBPD’s preferential regulation of PDGFRα involves BRD4

To further interpret the BRD4/CEBPD partnership at the next level, namely target gene expression downstream of CEBPD’s TF function, we were intrigued by literature evidence of their functional convergence. While our group reported a BRD4 preferential regulation of PDGFRα over PDGFRβ in SMCs^4^, another group reported that CEBPD regulated PDGFRα transcription preferentially over PDGFRβ in SMCs^22^. It is important to address the mechanism underlying this convergence as both PDGFRα and PDGFRβ are potent and key signaling mediators of multiple pro-IH SMC state transitions including SMC inflammation and proliferation/migration^4, 22^. As shown in Figure 7 (A-D), in our experiments using TNFα to stimulate the inflammatory SMC state transition, this treatment upregulated PDGFRα markedly yet PDGFRβ to a lesser degree especially at the protein level. Using JQ1 to block BRD4’s function abolished this upregulation. In parallel, CEBPD gain-of-function increased PDGFRα mRNA by ~2-3 fold in the presence or absence of TNFα but insignificantly affected PDGFRβ (Figure 7, E and F). CEBPD silencing abolished TNFα-stimulated PDGFRα upregulation (Figure 7, G and H). Of note, in accordance with the observed TNFα-stimulated BRD4 and CEBPD upregulation (Figure 6) and their function in preferentially up-regulating PDGFRα vs PDGFRβ (Figure 7), TNFα treatment alone preferentially elevated PDGFRα mRNA and protein levels with only a minor effect on PDGFRβ expression (Figure 7, A-H). As such, the CEBPD/BRD4 functional convergence contributes another layer of interpretation of their partnership from the angle of TNFα-stimulated CEBPD’s TF function entailing its direct target genes represented by PDGFRα.

**Figure 7.**
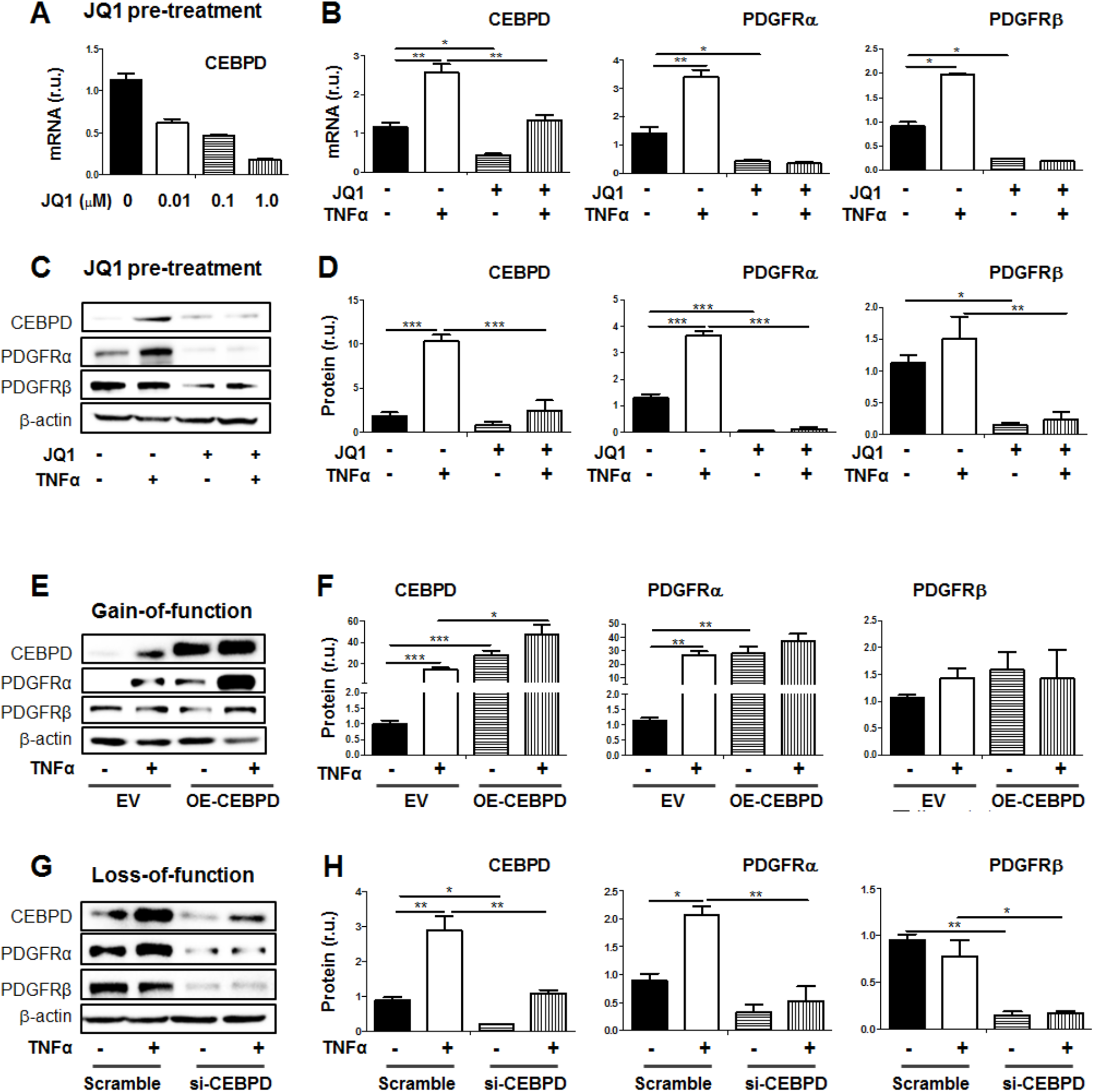
Preferential regulation of PDGFRα over PDGFRβ in SMCs. **A** - **D**. Effect of JQ1 pretreatment on TNFα-stimulated upregulation of CEBPD and PDGFRα. A shows concentration-dependent effect of JQ1 on reducing CEBPD mRNA levels. Representative Western blots are shown in C. **E** and **F**. CEBPD gain of function. Representative Western blots are shown in E. **G** and **H**. CEBPD loss of function. Representative Western blots are shown in G. MOVAS cells for stable expression of HA-tagged empty vector (EV) or HA-tagged CEBPD (OE-CEBPD) were starved in basal medium (0.5% FBS) for 24h, pretreated with vehicle or JQ1 (1 μM) for 2h, and then treated with TNFα (final 20 ng/ml) for 24h prior to harvest for qRT-PCR (mRNA) or Western blot (protein) analysis. For CEBPD silencing, MOVAS cells were transfected with siRNA for 24h and cultured in fresh starvation medium (0.5% FBS) for another 24h prior to TNFα treatment. Quantification: Readings from triplicate qRT-PCR reactions were normalized (to GAPDH) and averaged. The average values from at least 3 independent repeat experiments were then averaged again to calculate mean ± SEM (n =3-5). Densitometry of Western blots from independent repeat experiments was normalized (to β-actin, similar band intensity among blots) and then averaged to calculate mean ± SEM (n =4). Statistics: One-way analysis of variance (ANOVA) followed by Bonferroni post-hoc test; *P<0.05, **P<0.01, ***P<0.001.

**Figure 8.**
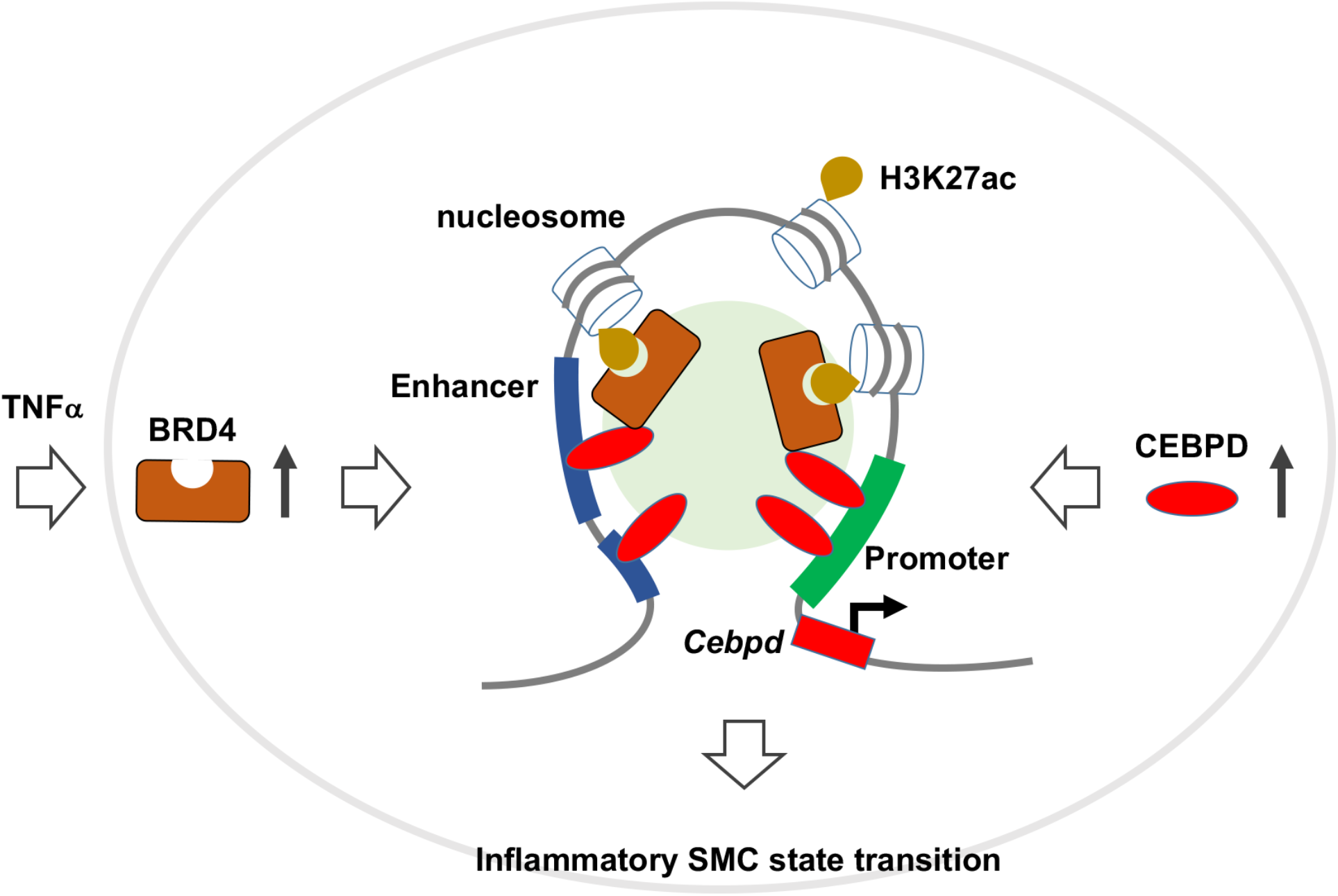
Schematic proposal for a BRD4/CEBPD partnership associated with chromatin. TNFα as extracellular signal stimulates upregulation of epigenetic reader protein BRD4 which, while reading H3K27ac, functions in the same complex with transcription factor CEBPD (and unknown co-factors, green circle in the back). BRD4 and CEBPD stock up at enhancers and *Cebpd* promoter to prompt the transcription of *Cebpd* (and other target genes); as a result, the inflammatory SMC state transition is propelled.

## Discussion

IH perpetuates vascular diseases. The pathogenic basis traces to SMC state transitions, whereby SMCs acquire new phenotypes (e.g. multiplied inflammatory cytokine expression) at the expense of losing their normal homeostasis and function. This SMC identity change is increasingly recognized as propelled by epigenetic remodeling^2^; thus, it becomes a compelling task to interpret the largely undefined mechanisms. We tackled this issue from the perspective of an interplay between BRD4, TF, and enhancer – key factors in cell state/identity changes^27^. Through ChIPseq genome-wide survey using arteries undergoing IH, we observed enrichment of BRD4 and H3K27ac at *Cebpd*. Genetically abating BRD4 (in vivo and in vitro) repressed *Cebpd* expression. Moreover, in the co-IP and ChIP-qPCR experiments, BRD4 and CEBPD were found to co-function in the same complex that also included CEBPD’s own promoter. Thus, these results revealed BRD4/CEBPD physical and functional associations. This BRD4/CEBPD partnership further manifested as promoting the inflammatory SMC state transition and their preferential transcriptional control of PDGFRα over PDGFRβ.

A unique interesting finding in this study is that CEBPD promotes its own transcription in SMCs, appearing to fit the concept of master TF. Among over a thousand TFs, a small number of so-called master TFs are critically important for cell identity, in other words, cell type or state^7, 28, 29^. Exemplary of master TFs, c-Myc is a powerful driver of oncogenic cell sate transitions^7^. In the current study, elevating CEBPD dramatically increased the expression of salient inflammatory SMC state markers including IL-1β, IL-6, and MCP1, and silencing CEBPD kept these markers at basal levels. Obviously, CEBPD plays a key role in the transition of SMCs to an inflammatory state. Moreover, master TFs are often found to associate with enhancers and BRD4 on chromatin^29, 30^; so is true for CEBPD as herein shown. First, ChIPseq data obtained from chromatin samples and bioinformatics identified enhancer regions enriched with BRD4 and H3K27ac. Second, a physical association between CEBPD and BRD4 was revealed by their co-IP. Third, ChIP-qPCR indicated that BRD4/H3K27ac-marked enhancer DNA was pulled down by ChIP against CEBPD. Interestingly, echoing our finding, CEBPD was recently found to bind inflammatory enhancers in aortic endothelial cells, although a BRD4 involvement was not reported^27^. Another prominent feature of master TFs is their regulation of own transcription, a mechanism thought to efficiently and rapidly activate transcription and cell state transitions in response to environmental perturbation^7^. Indeed, ChIP-qPCR indicated that IP against CEBPD was able to pull down its own promoter DNA, indicative of a physical association.

Essentially all the CEBPD physical associations, including that with enhancer DNA, its own promoter DNA, as well as that with BRD4 (co-IP), could be abolished by using JQ1 to block BRD4’s bromodomains. Importantly, aside from the CEBPD/BRD4 physical association, we also observed their functional association. First, BRD4 determined CEBPD expression levels; the role of BRD2 and BRD3 was minor (if any). We further distinguished that BD1 but not BD2 was critical for this BRD4 function – a finding worth more attention since BD1 and BD2 are highly similar and hence poorly differentiated in their biological functions^5, 19, 21^. Second, while TNFα as an extracellular stimulant upregulated CEBPD which prompted inflammatory SMC state transition (increased IL-1β, IL6, and MCP-1), using JQ1 to block BRD4 function abolished this TNFα stimulation. Third, CEBPD overexpression upregulated PDGFRα preferentially over PDGFRβ transcription, and TNFα stimulation had a similar effect which was abolished by JQ1. Apparently, the BRD4 and CEBPD functions converged. Moreover, indicative of a functional hierarchy of BRD4 and CEBPD, while TNFα upregulated both BRD4 and CEBPD, BRD4 governed CEBPD mRNA and protein production. Although JQ1 used to block BRD4 function also binds to the bromodomains of other BETs, we could attribute its inhibitory effect to BRD4 because our data established that BRD4 but not BRD2 or BRD3 dictated CEBPD expression, as iterated above.

Thus, we have identified a novel partnership between a powerful epigenetic reader (BRD4) and a master TF (CEBPD) which are physically and functionally associated in the complex that also includes BRD4-dependent enhancer and CEBPD’s own promoter. The function of this partnership manifested in the specific setting of TNFα-induced inflammatory SMC state transition, and in the transcription of *Cebpd* and CEBPD’s other target genes represented by PDGFRα, a signaling mediator crucial to SMC pathophysiology. As TFs bind to sequences of promoters and enhancers that are specific to the respective genes^27^, a TF/BRD4 association is thought to help localize BRD4 to certain genomic loci thereby defining its functional specificity. It is noted that this TF/BRD4 pairing is highly cell type and environment/stimulation dependent^6^. For instance, in the inflammatory endothelial cell state transition, NFκB was the master TF that paired with BRD4 to potently propagate this pathogenic process^29^. To the best of our knowledge, the BRD4/CEBPD partnership was not previously recognized, likely due to a paucity of information on CEBPD in contrast to extensively studied CEBPA. A BRD4 functional association with CEBPA in adipogenesis was recently reported^28, 30^, highlighting differential and contextual functions of different CEBP family members.

In the BRD4/CEBPD partnership observed here, BRD4 appears to play a central role orchestrating a multi-factor assembly that drives the inflammatory SMC state transition. As depicted in the schematic for the experimental setting herein applied, TNFα as extracellular signal stimulates upregulation of BRD4 protein which, while stocking up at enhancers and H3K27ac, partners with CEBPD to promote the transcription of *Cebpd* and other target genes, and the inflammatory SMC state transition results. This proposition is analogous to a model emerging from research in cancer and other fields^6, 7^, namely, BRD4 “rallies” enhancer-enriched TF(s) with the transcription elongation machinery and co-factors (e.g. MED1) while reading histone acetylation marks via its bromodomain(s)^2^. By so doing, BRD4 may “usher” the multi-factor assembly to select gene loci enabling their quick activation which drives cell state changes. As such, though seemingly a global regulator, BRD4 may assume functional specificity via context-specific associations with combinatorial TFs and enhancers at chromatin sites bookmarked (e.g. H3K27ac) in the epigenomic landscape. Nevertheless, since little is known about the BRD4/CEBPD duet in SMC (or other) cell state transitions, more research would generate exciting new knowledge to help decipher the underlying mechanisms.

## Conclusions

Our results reveal a previously unidentified BRD4/CEBPD physical and functional partnership that underlies the inflammatory SMC state transition, a process permissive for IH. A long-standing barrier in translational medicine is that TFs and enhancers inherently lack druggability limiting their targetability. Serendipitously, as implicated herein, their functional potency incumbent on BRD4 exposes an “Achilles’ heel”, that is, the druggable BRD4 bromodomains, targeting which could collapse the BRD4 assembly with context-specific TFs and enhancers. Therefore, justification is strong for more research on the BRD4/CEBPD partnership, so that essential information would become available for precision-oriented therapeutic interventions, in the vascular field or beyond.

## Methods

### Materials

Various resources including kits and reagents are presented in Table S1.

### Animals

All animal studies conform to the *Guide for the Care and Use of Laboratory Animals* (National Institutes of Health) and protocols have been approved by the Institutional Animal Care and Use Committee at The Ohio State University (Columbus, Ohio). Male Sprague-Dawley rats purchased from Charles River Laboratories (Wilmington, MA) were used for experiments (at 300-350 g of body weight).

### Rat carotid artery balloon angioplasty model

To induce IH, angioplasty was performed to injure rat carotid arteries, following our previous reports^4^. Briefly, rats were kept anesthetized with 2-2.5% isoflurane (inhaling, 2 L per min). After the left common carotid artery was dissected, a 2-F balloon catheter (Edwards Lifesciences, Irvine, CA) was inserted into the common carotid artery through an arteriotomy on the external carotid artery. The balloon was inflated (at 1.5 atm), withdrawn to the carotid bifurcation, and then deflated. This action was repeated three times. Perivascular BETs inhibitor administration was performed immediately following angioplasty as described below. Blood flow was finally resumed and the neck incision was closed. The animal was kept on a 37°C warm pad to recover. For postoperative analgesia, in addition to carprofen and bupivacaine, buprenorphine (0.03 mg/kg) was subcutaneously injected.

#### Artery tissue ChIP sequencing and data processing

To preserve the artery “real-time” epigenetic information, balloon-injured and uninjured (contralateral) common carotid arteries were snap frozen in liquid N_2_ immediately after dissected and severed out. Artery collection was performed 7 days after balloon angioplasty. Artery tissues from 40 rats were pooled for ChIPseq analysis at Active Motif per company standard procedures. Briefly, chromatin was isolated after adding lysis buffer, followed by disruption with a Dounce homogenizer. Genomic DNA was sheared to an average length of 300-500 bp by sonicating the lysates, and the segments of interest were immunoprecipitated using an antibody (4μg) against BRD4, H3K27a, H3K27me3, or H3K4me1. The protein/DNA complexes eluted from beads were treated with RNase and proteinase K, crosslink was reversed, and the ChIP DNA was then purified for use in the preparation of Illumina sequencing libraries. Standard steps included end-polishing, dA-addition, adaptor ligation, and PCR amplification. The DNA libraries were quantified and sequenced on Illumina’s NextSeq 500, as previously described^16^. Sequence reads were aligned to the reference genome Rn5, peak locations were identified using Macs2 algorithm^31^ and annotated based on UCSC RefSeq. Differential peak locations were called using SICER^32^. In-house shell and R scripts (https://www.r-project.org) were used for data integration. To summarize and cluster genome-wide TSS coverage as heat maps, deepTools (PMID: 24799436) compute matrix and plotheatmap functions were utilized. IGV (http://www.broadinstitute.org/igv/) was used for visualization. Annotation files were downloaded from UCSC.

### Immunofluorescence staining on artery cross sections

We followed our published protocol^33^. Briefly, artery sections were incubated without (negative staining) or with a primary antibody for 12 h and rinsed at least 3 times. The sections were then incubated with an anti-rabbit/mouse secondary antibody conjugated with Alexa Fluor 594 (A-11037/A-21203, Invitrogen, Carlsbad, CA) and rinsed. The specific antigen was then visualized with fluorescence microscopy. Detailed information of antibodies is included in Table S2. For quantification, 5 immunostained sections from each animal were used. Fluorescence intensity in each image field was quantified by using an ImageJ software and normalized to the number of DAPI-stained nuclei in the media and neointima layers. The values from all 5 sections were pooled to generate the mean for each animal. The means from all animals in each group were then averaged, and the final mean (±SEM) was calculated.

### Vascular smooth muscle cell (SMC) culture and transfection with siRNA

SMCs (MOVAS, mouse SMC line) were cultured in DMEM supplemented with 10% FBS and 100U/mL Penicillin-Streptomycin with 5% CO_2_ at 37°C. The construct for expressing dominant-negative BD1 of BRD4 was generated as we recently reported. For transfection with siRNA, cells cultured to 60%~80% confluence, then RNAi Max transfection reagent was added and incubated for 12h. The cells recovered in fresh DMEM (no Lipofectamine, 0.5% FBS) for 12h and starved for 24h in DMEM with 0.5% FBS. The cells were then incubated with 20 ng/ml TNFα or solvent control (0.1 BSA, 4mM HCl) for another 24h before harvest for various analyses. The siRNA sequences are listed in Table S3.

### Lentiviral constructs for expressing dominant-negative BRD4 bromodomains

Construction of vectors for the expression of GFP (control) or its fusion with BRD4-BD1 or BRD4-BD2, lentivirus packaging in Lenti-X 293 cells, and transduction of MOVAS cells were performed as we recently reported^5^ with minor modifications. The GFP lenti-vector was kindly provided by Dr. Ming-Liang Chu (Guizhou Renmin Hospital, China). Briefly, the crude viral solution was concentrated using Lenti-x concentrator (Takara, cat.631232) to a final concentration of 108-109 IFU/ml using the Lenti-x qRT-PCR Titration Kit (Takara, cat.631235). Lentivirus with MOI of 10 was used for transduction of MOVAS cells. Cells cultured to 70% confluency were changed to starvation medium (0.5% FBS), and lentivirus plus polybrene (Santa Cruz., Cat.sc-134220) was added and incubated for 6h. The cells were then cultured for recovery in fresh full medium (10% FBS) for 24h before use in experiments.

### SMC stable cell line for CEBPD overexpression

The CEBPD gene was cloned into the Lenti-HA-Vector (from Addgene). For lentivirus packaging, the construct of empty vector (Lenti-HA) or that for CEBPD overexpression (Lenti-HA-CEBPD) transfected into the LentiX-293 cell line together with plasmids PMD2G and PSPAX2. After a 24h culture, the super supernatant was collected. Lentivirus was purified and used to transduce the mouse smooth muscle cell line (MOVAS), and the stable cell line for the expression of HA control or HA-CEBPD was selected against puromycin.

### Western blot analysis

MOVAS cells were lysed in RIPA buffer. After quantifying with DC protein assay, equal amount of protein (20-40 μg) was loaded to and separated in 9% gel by SDS PAGE and then transferred to polyvinylidene difluoride (PVDF) membrane. The membrane was incubated with anti-PCNA or anti-actin overnight at 4°C, rinsed 3x, and then incubated with a peroxidase-conjugated secondary antibody for 1h at room temperature. Specific bands were illuminated by applying enhanced chemiluminescence (ECL) reagents (Thermo Fisher Scientific; Catalog no. 32106) and then recorded with Azur LAS-4000 Mini Imager (GE Healthcare Bio-Sciences, Piscataway, New Jersey). Band intensity was quantified with ImageJ. All antibodies are included in Table S2.

### Quantitative real-time PCR (qRT-PCR)

Total RNA was extracted from cell lysates using the TRIzol reagent following the manufacturer’s instruction (Thermo Fisher Scientific, 15596026) and used for cDNA synthesis with the High-Capacity cDNA Reverse Transcription kit (Thermo Fisher Scientific, 4368814). In each 20 μl reaction, 10 ng of cDNA was amplified through qRT-PCR using PowerUp SYBR Green Master Mix (Thermo Fisher Scientific, A25778), and mRNA levels were determined using 7500 Fast Real-Time PCR System (Applied Biosystems, Carlsbad, CA). The data was normalized to glyceraldehyde 3-phosphate dehydrogenase (GADPH) using the ΔΔCt method. The primers are listed in Table S4.

#### Co-immunoprecipitation (Co-IP)

We used Pierce Crosslink Immunoprecipitation Kit (Thermo Scientific, 26147) by following the manufacturer’s instruction. Briefly, cells were rinsed and incubated with ice-cold hypotonic buffer (2 Å~ 1 min, 20 mM HEPES, 20% glycerol, 10 mM NaCl, 1.5 mM MgCl2, 0.2 mM EDTA and 0.1% NP-40) supplemented with 1 mM dithiothreitol, and protease and phosphatase inhibitor cocktail (Thermo Fisher Scientific, 87785). Nuclei were collected and sonicated, and the lysates were cleared by centrifugation. Magnetic beads (Dynabeads Protein A or G, Invitrogen) preloaded with an anti-HA antibody were added to the supernatant and incubated at 4°C for 4h. The beads were washed 3x with the binding buffer (50mM Tris-Cl, 150mM NaCl, 1mM EDTA, 10% glycerin) and SDS sample buffer was then added to elute the co-IP’ed proteins for Western blot determination.

#### Chromatin immunoprecipitation (ChIP)-qPCR assay

ChIP was performed as we recently reported^34^ using the Pierce Magnetic ChIP kit (Thermo Fisher Scientific, 26157). Briefly, cells were cross-linked with 1% formaldehyde and the reaction was stopped by glycine. The cells were washed and lysed for nuclei extraction. Micrococcal nuclease was added to the nuclei suspension to digest the DNA for 15 min at 37°C, and then MNase Stop Solution was added to stop the reaction. The recovered nuclei were re-suspended in IP Dilution Buffer and sonicated (Four 5-second pulses at 20 Watts for 1×10^6 cells) to disrupt the nuclear membrane. Chromatin extracts containing DNA fragments (~500 base pairs in each) were immunoprecipitated by incubating with a specific antibody (or IgG) overnight at 4°C. ChIP-grade Protein A/G Magnetic beads were added and incubated for ~2-4h at 4°C. RNAse A and Proteinase K were used to digest RNA and protein. The purified DNA was used for qRT-PCR as described above. The primers are listed in Table S4.

### Statistical Analysis

Data are presented as mean ± standard error of the mean (SEM). Normality of the data was assessed based on Shapiro-Wilk normality test prior to statistical calculation using Prism 6.0 software (GraphPad). *p* < 0.05 was considered significant. One-way ANOVA followed by Bonferroni post-hoc test was applied to, as specified in each figure legend. For ChIPseq data, ***s***tatistical analyses were performed using SAS/STAT software, version 9.2 (SAS Institute, Inc., Cary, NC).

## Acknowledgement

This work was supported by NIH grants R01 HL133665 (to L.-W. G.), R01HL-143469 and R01HL-129785 (to K.C.K., L.-W. G.).

## Supplemental Tables

**Table S1.** Kits and reagents

| Product | Manufacturer | Catalog Number |
| --- | --- | --- |
| Protein A/G beads | Thermo Scientific | 88802 |
| RNAi Max | Invitrogen | 13778150 |
| Protease inhibitor cocktail | Thermo Scientific | 87785 |
| jetPRIME transfection reagent | VWR | 114-07 |
| PageRuler™ Plus Prestained Protein Ladder | Thermo Scientific | 26620 |
| Clarity Western ECL Substrate | Bio-rad | 170-5060 |
| QIAGEN Plasmid MIDI Kit | QIAGEN | 12143 |
| PureLink™ Quick Plasmid Miniprep Kit | Invitrogen | K210011 |

**Table S2.** Antibodies

| Antigen | Manufacturer | Catalog Number | Dilution Ratio | Application |
| --- | --- | --- | --- | --- |
| BRD2 | Proteintech | 22236-1-AP | 1:1000 | Western Blot |
| BRD3 | Proteintech | 11859-1-AP | 1:1000 | Western Blot |
| BRD4 | Abcam | ab128874 | 1:1000 | Western Blot |
| β-actin | Proteintech | 66009-1-Ig | 1:5000 | Western Blot |
| HA | Proteintech | 51064-2-AP | 1:5000 | IP |

**Table S3.**
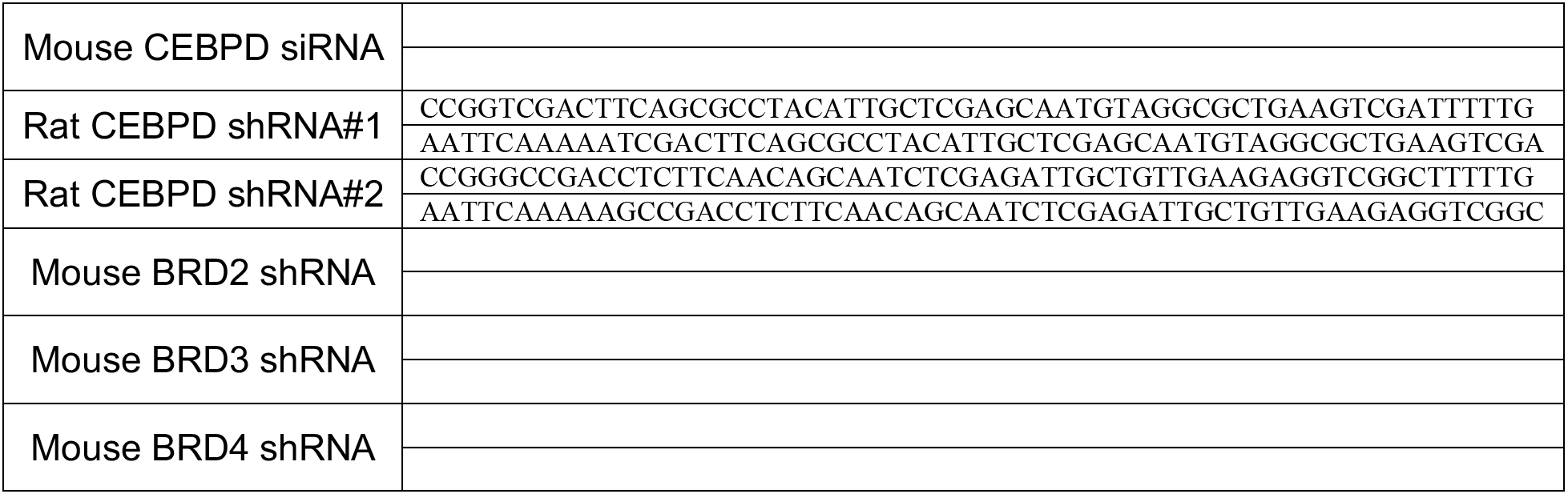
siRNA or shRNA sequences for mouse and rat genes

**Table S4.**
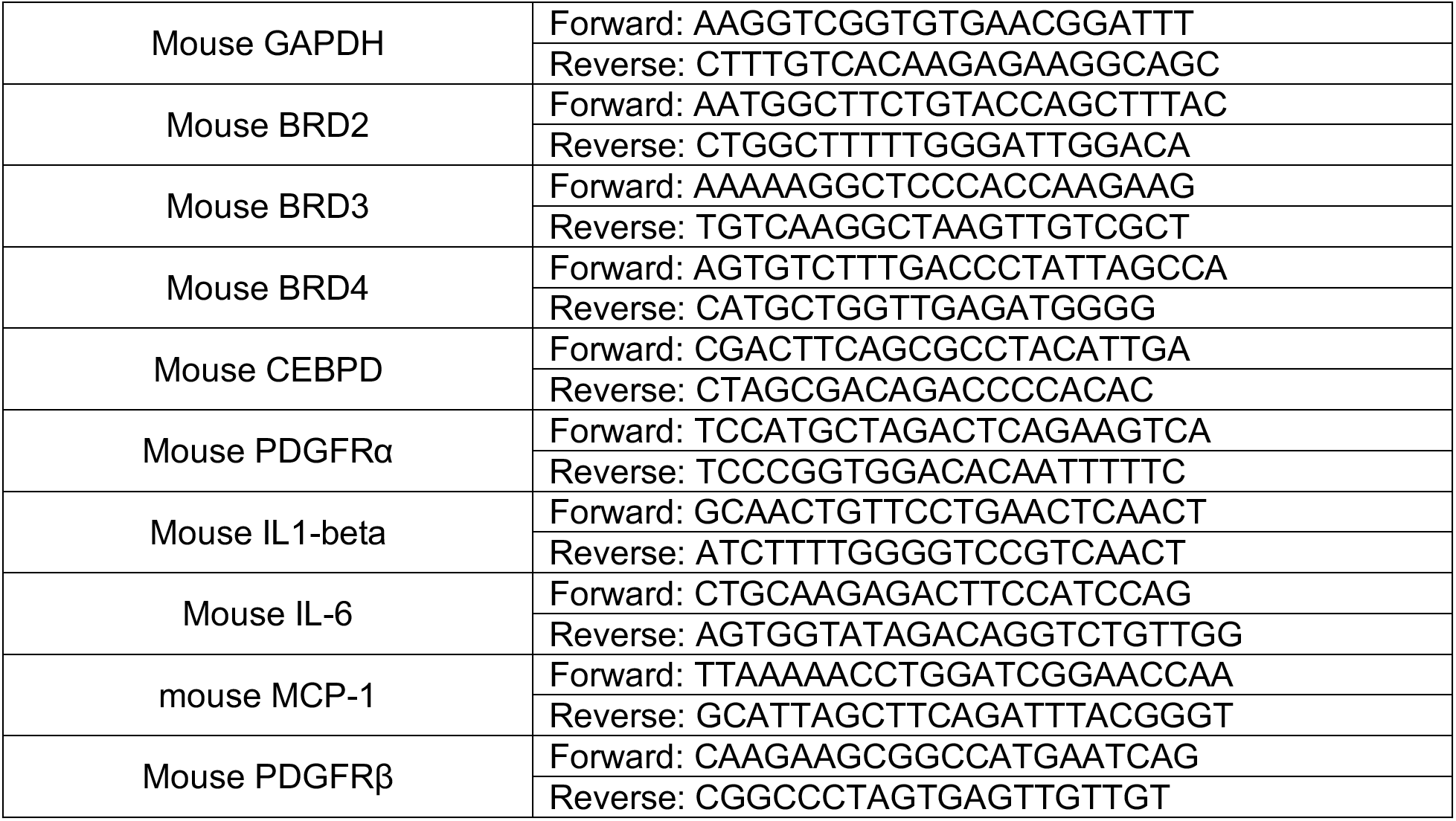
Primer sequences for mouse genes

